# Dynamic neuroinflammatory profiles predict Alzheimer’s disease pathology in microglia-containing cerebral organoids

**DOI:** 10.1101/2023.11.16.567220

**Authors:** Madison K. Kuhn, Rachel Y. Kang, ChaeMin Kim, Yerbol Tagay, Nathan Morris, Erdem D. Tabdanov, Irina A. Elcheva, Elizabeth A. Proctor

## Abstract

Neuroinflammation and the underlying dysregulated immune responses of microglia actively contribute to the progression and, likely, the initiation of Alzheimer’s disease (AD). Fine-tuned therapeutic modulation of immune dysfunction to ameliorate disease cannot be achieved without the characterization of diverse microglial states that initiate unique, and sometimes contradictory, immune responses that evolve over time in chronic inflammatory environments. Because of the functional differences between human and murine microglia, untangling distinct, disease-relevant reactive states and their corresponding effects on pathology or neuronal health may not be possible without the use of human cells. In order to profile shifting microglial states in early AD and identify microglia-specific drivers of disease, we differentiated human induced pluripotent stem cells (iPSCs) carrying a familial AD PSEN2 mutation or its isogenic control into cerebral organoids and quantified the changes in cytokine concentrations over time with Luminex XMAP technology. We used partial least squares (PLS) modeling to build cytokine signatures predictive of disease and age to identify key differential patterns of cytokine expression that inform the overall organoid immune milieu and quantified the corresponding changes in protein pathology. AD organoids exhibited an overall reduction in cytokine secretion after an initial amplified immune response. We demonstrate that reduced synapse density observed in the AD organoids is prevented with microglial depletion. Crucially, these differential effects of dysregulated immune signaling occurred without the accumulation of pathological proteins. In this study, we used microglia-containing AD organoids to quantitatively characterize an evolving immune milieu, made up of a diverse of collection of activation patterns and immune responses, to identify how a dynamic, overall neuroinflammatory state negatively impacts neuronal health and the cell-specific contribution of microglia.

## Introduction

The multifaceted disease-driving mechanisms that underlie Alzheimer’s disease (AD) have complicated the development of successful therapeutic interventions. Whether the disease is familial (inherited through disease-causing genetic mutations) or sporadic (initiated by a combination of genetic, lifestyle, and environmental risk factors) in origin, it is characterized by the largely predictable accumulation and spread of amyloid beta (Aβ) plaques and neurofibrillary tangles of primarily hyperphosphorylated tau (pTau) in the brain. However, the pathological events in the initiation of both types of AD are not fully understood. Neuroinflammation is an early attribute of the AD brain and can actively worsen disease through numerous aberrant immune responses including the secretion of cytokines that directly injury neurons, hyper-phosphorylate tau, or cause the production of reactive oxygen species.^1,2^ Microglia, the resident immune-competent cells of the brain, are the primary cells driving neuroinflammation,^3^ and likely play a critical role in early disease, as revealed by the numerous genetic risk factors that are uniquely expressed by microglia,^4,5^ the ability of reactive microglia to initiate neurodegeneration,^6^ and microglia-dependent mechanisms driving proteinopathy.^3,7^ Although correcting their dysfunction may throw disease off its course, characterizing diverse microglial states has proven challenging, as unique immune stimuli and situational- or temporal-dependent cues may produce drastically different, and sometimes contradictory, responses.^3,8^ These activation states cannot be easily teased apart, but transcriptomic studies have demonstrated their large diversity and dynamic nature, identifying a predictable evolution of states.^9–11^ How transcriptomic signatures translate to microglial dysfunction, however, is difficult to characterize and is made more challenging because of the differences between human and mouse microglia and AD models.

Mice and human microglia have critical differences that obscure physiological relevance, such as opposing gene regulation,^12,13^ human-specific activation patterns, and decreased neuroinflammation in AD mouse models compared to the human brain.^14^ Additionally, mice do not naturally develop AD and the overexpression of familial AD mutations is often used to produce AD pathology in the mouse brain. However, pathology is often incomplete^14,15^ and accompanied by confounding effects of gene overexpression and the interaction of humanized proteins with mouse proteins and microenvironment.^15^ Importantly, these factors may completely misrepresent events that initiate disease. Human induced pluripotent stem cell (iPSC)-derived models are becoming more widely used in AD research and are appreciated for their ability to more accurately represent Aβ and tau pathology in the context of the human genome, which can be derived from both familial and sporadic patients.^16^ Cerebral organoid models further improve cell-cell complexity and the development of more AD-reminiscent features.^17–19^ Recently, microglia have been introduced into organoid models, either by their separate introduction or fostering their innate development. ^20–22^ Despite the growing appreciation for the importance of microglia in AD and human-specific responses, microglia-containing cerebral organoids have not yet been investigated in the study of AD. In this study, we employ a previously developed cerebral organoid model^23^ that was optimized to give rise to microglia^22^ to investigate microglia-driven changes and characterize the corresponding overall neuroinflammatory state.

The hiPSCs used in the study for AD modelling carry the familial presenilin 2 (PSEN2) mutation N141I, that predominantly causes an overall heighten neuroinflammatory state with exaggerated glial immune responses in the presence of immune insult, such as Aβ.^24,25^ The exaggerated immune response accompanied by the N141I mutation is reminiscent of a “primed” microglial phenotype^26^ that describes an over-reactive state with amplified cytokine secretion as a result of a chronic inflammatory environment commonly observed with age and AD.^27^ In order to characterize changing immune responses in the organoid model over time, we profiled cytokine secretion in organoids derived from a male hiPSC line carry the PSEN2 N141I mutation and its isogenic control and built cytokine signatures predictive of age and genotype using the multivariate modeling tool partial least square (PLS) to identify key cytokines and differential regulation that correlated with observed pathology changes in the organoids. We identified an initially amplified cytokine profile that gradually decreased over months in culture, which coincided with decreased synapse density with minimal changes in the levels of pathological AD proteins. Microglial depletion prevented synapse loss and supports previously identified pathology-independent microglia-driven neuronal injury.

## Methods

### iPSC culture

We purchased previously characterized human iPSC lines for the generation of cerebral organoids. The iPSC line JIPSC1052 (SNV/WT, Jackson Laboratory), carrying the familial AD PSEN2 mutation N141I, and its isogenic control JIPSC1054 (REV/WT, Jackson Laboratory) are CRISPR-edited cell lines of an original parent iPSC line derived from a white male of 55-59 years of age (KOLF2.1J).^28^ The isogenic control is a CRISPR-edited reversion of the CRISPR-introduced PSEN2 mutated gene back to wild-type to serve as a more robust control, having also gone through the gene editing process. iPSCs were maintained in mTeSR1 medium (STEMCELL Technologies) on Matrigel-coated plates (Corning, cat. 356234) in a 5% CO2, 37°C cell culture incubator with daily medium changes. iPSC cultures were passaged every 4-5 days when the colonies reached 80% confluency using the dissociation reagent ReLeSR (STEMCELL Technologies) according to the manufacturer’s protocol. iPSCs were passaged 5-6 times post thaw prior to starting organoid differentiation. iPSC cultures were monitored daily for appropriate morphology and mycoplasma detection was performed every two-weeks in culture (MycoAlert, Lonza LT07-318).

### Organoid differentiation and maintenance

The male N141I PSEN2 AD iPSC line (JIPSC1052, Jackson Laboratory) and its isogenic control (JIPSC1055, Jackson Laboratory) were differentiated into cerebral organoids according to a previously published protocol^23^ with minor modifications to support the differentiation of the specific iPSC lines and to optimize the organoids’ development of microglia.^22^ When iPSC cultures reached 80% confluency, the cells were dissociated into a single-cell solution with Accutase (STEMCELL Technologies), and 9,000 live cells were seeded in each well of U-bottom Ultralow attachment 96-well plates (Corning, cat. 7007) in mTeSR1 medium containing 10 μM Y-27632 ROCK inhibitor (STEMCELL Technologies) for the formation of embryoid bodies. iPSCs used in the generation of cerebral organoids demonstrated above 95% viability (Countess Automated Cell Counter, Invitrogen). After seeding the iPSCs in U-bottom plates (Day 1), a half change of the mTeSR1 medium was performed (without ROCK inhibitor) on Day 3. On day 5, the embryoid bodies were transferred to individual wells of Ultralow attachment 24 well plates (Corning, cat. 3473) using a cut p200 pipette tip. Embryoid bodies were cultured in the 24 well plates containing neural induction medium for 7 full days with medium changes on days 7 and 9. On day 12, the organoids were embedded in Matrigel and transferred to 60mm dishes containing cerebral organoid differentiation (COD) medium without vitamin A. Briefly, individual organoids were transferred to dimples in a Parafilm sheet with a cut p200 pipette tip (16 organoids/dish). Residual medium around each organoid was removed and 20-30μL of Matrigel was added drop-wise to each organoid dimple. Organoids were positioned to the center of the droplet using a p10 pipette tip. Each sheet of Matrigel-embedded organoids was incubated for 20 to 30 minutes at 37°C. After Matrigel polymerization, COD medium was added to the dish overtop the organoids to dislodge them from the Parafilm sheet, which was then removed. On day 14, the COD medium without vitamin A was replaced. On day 17, the medium was replaced with COD medium with vitamin A, and the dishes were placed on an orbital shaker at 80 rpm (KS 260 Control, IKA). Dishes were placed on the orbital shakers in stacks of 4 with 2 empty dishes on the bottom of each stack to prevent condensation from accumulating on the underside of the lids. Medium was replaced every 3-4 days. After 1 month in culture, the organoids were split into new dishes with 8 organoids per dish to prevent organoids from fusing together.^29^ Neural induction medium and COD medium was prepared according to the original protocol,^23^ except for the inclusion of 0.1μg/mL heparin, as opposed to of 1μg/mL, in the neural induction medium as detailed by Ormel *et al.*^22^ Large fused organoids or those not exhibiting optimal morphology^23,29^ were excluded from the study. All the organoids used in the study came from a single batch, denoted as the organoids produced from one single-cell solution preparation that underwent the same differentiation steps. Organoids were cultured for 2-, 3-, or 4-months prior to sample collection.

### PLX5622 treatment of organoids

For microglia depletion studies, PLX5622 (in DMSO, MedChemExpress) was added to the COD medium for a final concentration of 5 μM with each medium change. The volume of DMSO in the cell culture medium was less than 1%. PLX5622 or equal amounts of vehicle control was applied from 2- to 3-months, 3- to 4-months, or 2- to 4-months in culture prior to sample collection.

### Organoid preparation for immunostaining

For the fixation of organoids for immunostaining, groups of organoids were transferred to conical tubes and washed with 1X PBS. The PBS was replaced with 4% (wt/vol) PFA, and the organoids were incubated at room temperature for 15 minutes. The PFA was then aspirated, and the organoids were washed 3 times with 1X PBS, with a 10-minute gentle agitation on a rocker at room temperature for each wash. The final PBS wash was replaced with 30% (wt/vol) sucrose solution. Organoids in sucrose were incubated at 4°C on a rocker at very slow speed until all the organoids had sunk to the bottom of the tube (typically 48-72 hours). After the sucrose sink, organoids were transferred to embedding molds in OCT compound (Tissue-Tek). Embedding molds were filled hallway with OCT compound and frozen at −20°C. In each tube, half of the sucrose solution was removed and replaced with OCT compound. The organoids in sucrose/OCT were then incubated at room temperature for 30-60 minutes with gentle agitation. All of the sucrose/OCT solution was then replaced with OCT compound, and the tubes were incubated for another 30-60 minutes with gentle agitation at room temperature. After the equilibration with OCT, a thin layer of OCT was applied to the frozen OCT in the embedding mold. The organoids were then quickly transferred on top on the liquid OCT layer using a cut P1000 pipette tip (6 organoids per mold). Organoids were gently arranged as desired in the mold and more OCT was applied to cover them. The molds were quickly frozen on a metal block in liquid nitrogen. Frozen OCT-embedded organoids molds were stored at −80°C. Organoid blocks were cryosectioned using a cryostat (Leica) at 20μm at −20°C on Superfrost Plus Microscope Slides (Fisherbrand). Organoid slides dried completely at room temperature before being stored at −80°C.

For immunostaining, organoid slides were thawed and allowed to dry completely at room temperature before washing and blocking. Slides were washed in 1X PBS three times with gentle agitation for 5-10 minutes at room temperature. A hydrophobic barrier was drawn around each grouping of organoids (Super HT Pap Pen, Kiyota International) and blocking solution was added (90% PBS, 10% normal goat serum, and 1% of 10% Triton-X-100 Solution). Slides were incubated in the blocking solution at room temperature for 1 hour. The blocking solution was then replaced with primary antibodies in blocking solution (anti-TUBB3 (Abcam ab78078, ab52623), -AT8 (Thermo MN1020), - IBA1 (Wako 019-19741), -Tmem119 (Invitrogen PA5-119902), -GFAP (Abcam ab207165), SYN1 (Abcam ab254349), and -Aβ-40/42 (Millipore AB5076)). Slides were incubated at 4°C overnight in a humidity chamber. Slides were then washed with PBS 3 times and secondary antibodies (Alexa Fluor 555 goat anti-mouse (Invitrogen A21425) and Alexa Fluor 488 goat anti-rabbit (Invitrogen, A11070)) in blocking solution were added to the slides for 1 hour at room temperature. Following the secondary antibody incubation, slides were washed in a weak serum solution (1% normal goat serum, 99% PBS) for ten minutes with gentle agitation at room temperature and subsequentially washed an additional 2 times with PBS. Coverslips were mounted with Prolong Diamond Antifade Mountant with DAPI (Invitrogen) and placed flat in the dark for 24 hours at room temperature prior to storage at 4°C.

### Organoid imaging

Organoid slides were viewed and imaged using a fluorescence microscope (Nikon Eclipse 80i, Nikon DS-Fi3 camera, NIS-Elements AR software). Some fluorescent signals were not uniform across the organoid sections, such as the isolated clusters of astrocyte GFAP staining. In these instances, when the magnification did not allow for the entire organoid section to be imaged, images were collected from the region(s) of each organoid that displayed the largest signal intensity in order to accurately assess differences in pathology staining/ cell populations. Images of sections from the middle of the organoid displaying necrosis were excluded from analyses. Images were processed and analyzed in ImageJ. Background correction of single-color channel images was performed by subtracting the mean intensity of the background from image. Histogram stretching was performed on images not used for quantitative analysis; however, a minimum maximum value of 200 was used to prevent one color signal from over-powering the others. For the measurement of mean intensity of the organoid sections, outlines were manually drawn around each organoid and the mean intensity was measured. Large tears and ventricle-like pockets^29^ within the organoids were excluded from the mean intensity measurement. If the signal was too weak to see the outline of the organoid (as was the case in the

Tmem119 signal after microglia depletion), the brightness of the image was increased to draw the outline and was reset prior to taking the intensity measurement. Tmem119, rather than IBA1, was used for the PLX-treatment experiments to assess the depletion of microglia because it was more representative of the population within the whole organoid, as the expression was more widespread and consistent between sections of the same organoid. For Tmem119 signal intensity quantification, images were taken of whole organoids (4X magnification) to appreciate entire the population visible in a cross section, while Aβ and pTau images were taken at 10X magnification of organoid region(s) demonstrating the largest fluorescence intensity to better visualize the morphology of Aβ and pTau deposits.

For the quantification of Aβ deposits, the percentage of pixels contained within Aβ signal-rich regions was measured. First, a fluorescence intensity threshold value of 73 was determined by taking the average of the mean fluorescence intensity measurements of the largest Aβ deposits of the AD and isogenic organoid images. The background of each image around the organoid and any distinguishable holes in the organoid were removed (fluorescence intensity = 0). Each image was converted to a text file in ImageJ containing the fluorescence intensity of each pixel. In R, each pixel was assigned “0” for background, “TRUE” if it and its neighboring pixels were above 73, or “FALSE” otherwise. Pixels in the middle of the section had 8 neighbors, pixels on the edge had 5, and pixels in the corners had 3 neighbors. The percentage of “TRUE” observations over the total of “TRUE” and “FALSE” observations was recorded to quantitatively describe the concentration/diffusivity of the Aβ signal between organoids (Supplementary Figure 1).

### Organoid lysate preparation

Organoid lysate samples were collected for the quantification of cytokine and pathological protein concentrations in the organoids. Groups of 3 to 5 organoids were transferred to Eppendorf tubes and washed with cold 1X PBS. Organoids were combined for individual samples to have adequate total protein concentration and volume for multiple multiplex assays. To remove the Matrigel surrounding the organoids, the PBS was replaced with Cell Recovery Solution (Corning), and the organoids were incubated at 4°C for 30 minutes. The organoids were then washed with cold PBS. The tubes were briefly spun down and the PBS was aspirated. Cold cell lysis buffer (Millipore 43-040) containing protease inhibitor cocktail (1:100, Sigma) was added to the organoids, and the organoids were mechanically digested with trituration using a p1000 and then a p200 pipette. Following the trituration, the tubes were vortexed for 1-2 minutes and kept on ice at least 20 minutes. The samples were then centrifuged at room temperature for 10 minutes at 10,000 x g. The supernatants were transferred to a new tube and aliquoted for separate assays and BCA total protein quantification. The aliquots were flash frozen in liquid nitrogen and stored at −80°C. The total protein content of the organoid lysate samples was quantified using the Pierce BCA Protein Assay Kit (Fisher 23225) according to the manufacturer’s instructions. Each sample was run in triplicate and absorbances read using a SpectraMax i3 minimax 300 imaging cytometer (Molecular Devices). Sample concentrations were quantified by linear regression using triplicate standard samples ran on each plate.

### Luminex multiplex assays

The cytokine concentrations of organoid lysate samples were quantified on the Luminex FLEXMAP3D platform using the Milliplex human cytokine and chemokine magnetic bead panel kit (Millipore HCYTOMAG-60K), measuring a broad panel of immune signaling proteins that allow for an unbiased survey of immune cues activating a diverse set of downstream intracellular pathways. Aβ and pTau levels in the organoids were quantified with Milliplex human amyloid beta and tau kit (Millipore HNABTMAG-68K) on the Luminex. The assays were performed according to the manufacturer’s protocols with minor modifications to accommodate the use of 384-well plates. The magnetic beads and antibody solutions were diluted 1:1 and used at half volume, and the streptavidin-phycoerythrin was used at half volume. Samples were diluted to 1.2 mg/mL total protein using assay buffer and added to the plate for 30 μg total protein per well. Samples were assayed in technical triplicate.

### Cytokine profile data cleaning

Cytokine concentrations for each sample were interpolated from 5-point logistic standard curves using the Luminex Xponent software. Concentrations below the detection limit (< 3.2pg/mL) were assigned 0 pg/mL. The raw concentration data was processed using an automated in-house pipeline, available from GitHub at https://github.com/elizabethproctor/Luminex-Data-Cleaning (Version 1.02). The pipeline removes readings generated from less than 35 beads (custom input value) (0 observations in the datasets) and then calculates the pairwise differences of the remaining technical triplicates. If the difference between one replicate is greater than twice the distance between the other two, the replicate is removed from the dataset. The average of the remaining technical replicates for each cytokine is then calculated for the final dataset. We then manually removed entire cytokines from the datasets for subsequent analyses if over half the readings were 0 pg/mL but did not partition between experimental groups, such as genotype or timepoint.

### Partial least squares modeling

The linear, supervised multivariate mathematical modeling tool partial least squares (PLS)^30,31^ was used to construct cytokine signatures predictive of a response of interest (ex: AD vs control). Cytokine expression and signaling is highly interdependent, and PLS allows for the identification of significant multivariate changes in correlative predictors (the cytokines) as they relate to a dependent response or group (such as, genotype or timepoint). This is achieved through PLS’s construction of linear combinations of the predictors (latent variables, LVs) that maximize the covariation between the predictors and the response. This maximization of multivariate covariance in predictors with the response allows us to identify subtle but meaningful patterns in cytokine expression that are undetectable using univariate analysis methods, which are unsuitable for highly correlative variables. We generated PLS discriminant analysis (PLS-DA) models for the prediction of experimental groups using our previously published workflow^32^ in R with the *ropls* package.^33^ For each PLS-DA model, leave-one-out cross validation (n=10) was used to estimate the average classification error and determine the optimal number of LVs for each model. The cytokine data were mean-centered and unit-variance scaled prior to PLS modeling. The models were orthogonalized to maximally project covariation of the measured cytokines with the response to the first latent variable, which improves interpretability by prioritizing predictor/response covariation over variation in the measured cytokines between samples.^34^ A model’s significance was evaluated using a permutation test. In each iteration, the sample identities are randomly reassigned to unchanged cytokine signatures. The randomized data is used to generate models for cross validation to measure the performance of the random models compared to the “real” experimental model. The p-value of the optimized experimental model is then calculated by comparison of its accuracy with the mean and standard deviation of the distribution of random models’ accuracy. The VIP score, a measure of a variable’s normalized contribution to the predictive accuracy of the model across all LVs,^35^ was used to identify key cytokines in the models, where a VIP score > 1 indicates a greater than average contribution to the model.

### Single-Cell RNA Sequencing

We used the SPLiT-seq technique as commercialized by Parse Biosciences to conduct combinatorial barcoding of gene transcripts in single cells from our organoids, according to the manufacturer’s protocol. In brief, we dissociated organoids using papain and fixed cells using Parse Bioscience’s cell fixation kit. We prepared libraries using Parse Bioscience’s Evercode WT Mini whole transcriptome kit. We sequenced libraries on the Illumina NovaSeq sequencer (SP flow cell) to a maximum depth of 75,000 reads per cell.

### Statistical Analyses

Statistical tests were performed using Graph Pad Prism Version 10.0.1. To evaluate the statistical significance of changes in protein level or fluorescence intensity between genotype and timepoints, two-way ANOVA with Tukey’s multiple comparison test was used. A one-way ANOVA with Tukey’s multiple comparison test was used for timepoint comparisons in the PLX5622 and vehicle treatment comparisons. Unpaired two-tailed Student’s t-tests were used to determine significance between two populations. A Grubbs’ test was used to identify outliers in the datasets. All data are expressed as mean ± standard deviation.

## Results

### AD N141I and isogenic control iPSC-derived cerebral organoids innately develop microglia

A chronic, aberrant neuroinflammatory state contributes to Alzheimer’s disease proteinopathy accumulation and neuronal injury.^36^ The familial AD N141I PSEN2 mutation is observed to heighten the immune response of microglia and astrocytes.^26^ In order to dissect N141I-driven changes in glial function and the presentation of AD pathology, a hiPSC line carrying the familial AD PSEN2 N141I mutation and its isogenic control were used to generate cerebral organoids, which were cultured for 2-, 3-, or 4-months. It has been previously demonstrated microglia are found throughout the organoid after 2 months in culture and their transcriptomes closely resemble those of adult human microglia.^22^ The microglia continue to mature, increasing expression levels of microglia-specific genes, in the following months.^22^ In a pilot study, AD organoids were cultured for up to 6-months, and widespread tissue atrophy was observed by 5-months (Supplementary Fig 2). We chose to end the experimental timeline at 4-months to eliminate confounding effects of robust cell death on the immune response and pathology changes in the organoids. We first verified the existence of microglia and astrocytes in our organoids with the immunostaining of the cell-type specific markers GFAP (astrocytes) and Tmem119 and IBA1 (microglia)(Fig 1). Tmem119, a transmembrane protein expressed by microglia and not by other brain cell-types or infiltrating macrophages,^37^ is found throughout the organoids (Fig 1A). The Tmem119 signal is distinct from but closely associates with neuron projections (Fig 1B). The single cell preparation of the organoids demonstrates Tmem119-positive cells (Fig 1D). The organoids demonstrate clusters of astrocyte populations, which also closely associate with neurons (Fig 1C,E). The microglia activation marker, IBA1, is also expressed in the organoids, though much more sparsely (Fig 1E). Microglia and astrocyte populations overlap in the organoids (Fig 1E). It is worth recognizing that in the literature these cells derived directly from iPSCs or within organoids may not be fully mature microglia and are occasionally instead referred to as “microglia-like cells”.^22,38–42^ With the presence of Tmem119 and IBA1 positive cells, proteins regarded as microglia-specific markers in the brain,^22,37^ they will be referred to as microglia with the acknowledgement that their full likeness to endogenous human microglia is unknown.

**Figure 1:**
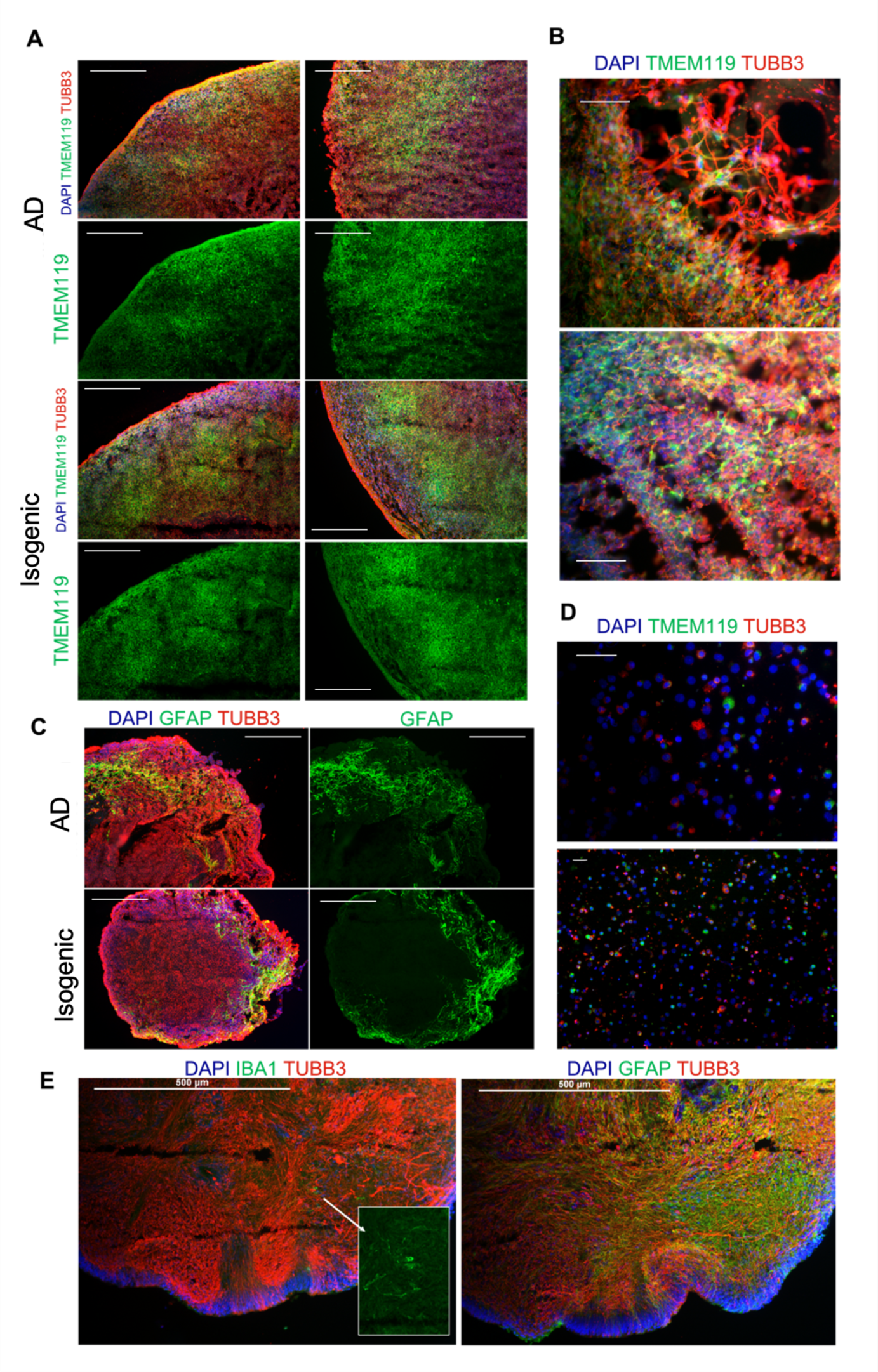
Microglia and astrocytes develop in AD and isogenic organoids. (A) Composite (red: TUBB3, green: Tmem119, blue: DAPI) and green channel (Tmem119) images of 3-month-old AD (top) and isogenic organoids (bottom), scale bars: 250μm. (B) Magnification of Tmem119 and TUBB3 signals emphasizes separate microglia and neuron projections, scale bars: 50μm. (C) Composite (red: TUBB3, green: GFAP, blue: DAPI) and green channel (GFAP) images of 3-month-old AD (top) and isogenic (bottom), scale bars: 250μm. (D) Tmem119 (green), TUBB3 (red), and DAPI (blue) immunostaining of a single-cell preparation of an isogenic control organoid, scale bars: 50μm. (E) Sequential organoid sections demonstrating overlapping microglia (IBA1, left) and astrocyte (GFAP, right) populations, scale bars: 500μm.

### Cytokine secretion is initially increased in AD organoids prior to an overall down-regulated cytokine profile compared to isogenic control organoids

To characterize the effect of the PSEN2 N141I mutation on the organoids’ inflammatory state, we profiled the immune milieu of AD and isogenic organoids by measuring cytokine concentrations over 2- to 4-months in culture. We used partial least squares discriminant analysis (PLS-DA) to construct cytokine signatures predictive of genotype (AD vs control) to identify changes in immune signaling in the organoids over time and distinguish mutation-dependent differences in the cytokine profiles (Fig 2). In 2-month-old organoids (Fig 2A. PLS-DA: 2 latent variables, accuracy: 90%, p-value: DAPI GFAP TUBB3 <0.005), unique cytokines are up-regulated in AD or control organoids. Key cytokines, determined by a VIP score greater than 1 (having a greater than average contribution to the model), up-regulated in the AD organoids are IL-4, IL-12p70, IFNα2, GRO, Flt-3L, and FGF2, while increased IP-10 (CXCL10) and IL-1Ra expression are key cytokines in the prediction of isogenic organoids. In 3-month-old organoids (Fig 2B. PLS-DA: 2 latent variables, accuracy: 80%, p-value: <0.05) there is a shift in the cytokine profile from that of the 2-month-old organoids. In the AD organoids, there is diminished expression of key 2-month AD cytokines, such as IL-12p70, IFNα2, GRO, and Flt-3L, and now undetectable levels of IL-4 (which was excluded from the model), while demonstrating an increase in VEGF, Fractalkine, and FGF-2. The cytokine signature of the AD 4-month-old organoids (Fig 2C. PLS-DA: 3 latent variables, accuracy: 80%, p-value: <0.05) demonstrates further reduction of all cytokines, excluding VEGF, showing a complete switch of the 3-month-old correlation of AD organoids with up-regulated Fractalkine and FGF-2. The comparison of cytokine signatures over the months in culture demonstrates an initial up-regulation of cytokines in AD organoids, which diminishes overtime and is completely reversed at 4-months where cytokines are up-regulated in the control organoids. PLS regression clearly demonstrates the overall reduction of cytokine secretion in AD organoids between 2 and 4 months (Supplementary Figure 3). Importantly a significant PLS regression model could not be generated for the cytokine profiles of the isogenic organoids, indicating there was no predictable signature of cytokines that differentiated increasing age in the isogenic controls.

**Figure 2:**
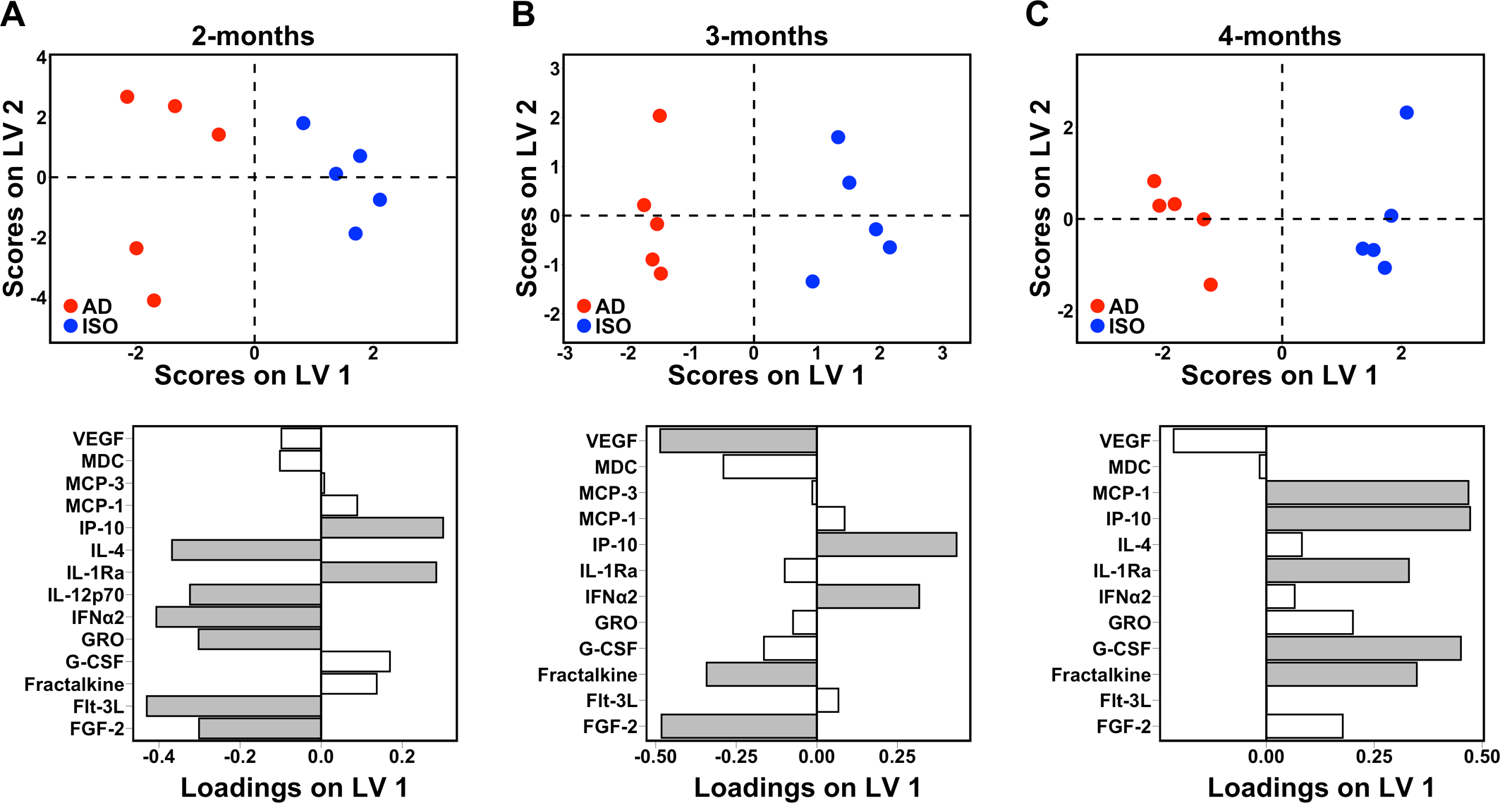
Cytokine signature predictive of genotype in AD organoids reverses over experimental timeline. PLS-DA scores plot (top) and LV1 loadings plot (bottom) of (A) 2-month-old AD vs isogenic control organoids (2 LV, accuracy: 90%, p-value: <0.005), (B) 3-month-old AD vs isogenic control organoids (2 LV, accuracy: 80%, p-value: <0.05), (C) 4-month-old AD vs isogenic control organoids (3 LV, accuracy: 80%, p-value: <0.05). Shaded loadings indicate a VIP score > 1. Each point represents a single sample comprising 3-5 organoids. Positive loadings indicate an up-regulation of cytokines with positive scoring samples, while negative loadings feature the down-regulation of cytokines with samples of positive scores.

### Down-regulation of cytokine secretion in AD 4-month-old organoids coincides with decreased synapse density

To verify that the decrease in cytokine secretion of aging AD organoids represented a diminished reactive glial phenotype and was not a result of cell death within the organoids, we confirmed that the organoids did not display tissue atrophy over the culture period (Fig 3A). Subsequently, we investigated if there was a change in the microglia population, as microglia-specific loss could also mimic the phenotype. We did not observe a loss of Tmem119 signal over the culture period in either the AD or isogenic organoids and instead saw an overall increase in signal from 2- to 4-months (Fig 3C). Thus, the decreased cytokine profile of aging AD organoids is not due to a loss of microglia over time and is likely a result of a dampened immune response. Notably, from 2- to 4-months, a reduction of SYN1, a presynaptic marker, coincided with the increase of the Tmem119 signal in the AD organoids (Fig 3B), indicating decreased synapse density. Initially at 2-months, there was increased SYN1 in AD organoids compared to controls (Fig 3B).

**Figure 3:**
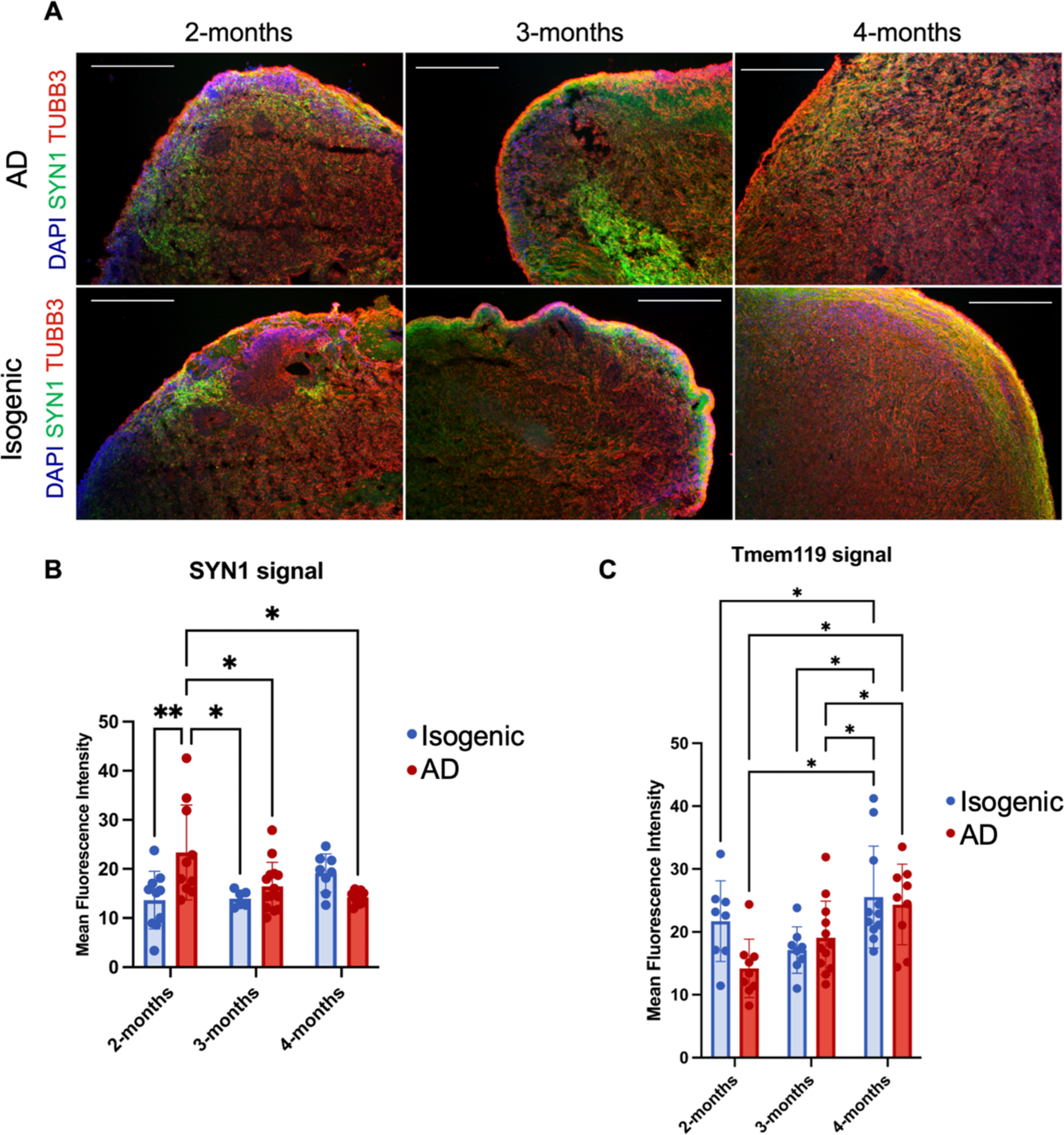
Synapse density decreases over time in AD organoids along with increase in microglia population. (A) Composite images (red: TUBB3, green: SYN1, blue: DAPI) of 2-, 3-, and 4-month-old AD (top) and isogenic organoids(bottom), scale bars: 250μm. (B) Quantification of mean fluorescence intensity of SYN1 immunostaining in organoid sections from 2- to 4-months. Two-way ANOVA with Tukey’s multiple comparison test. 2-month-old isogenic vs 2-month-old AD, p-value < 0.005. 2-month-old AD vs 3-month-old isogenic, p-value < 0.05. 2-month-old AD vs 3-month-old AD, p-value < 0.05. 2-month-old AD vs 4-month-old AD, p-value = 0.01. (C) Quantification of mean fluorescence intensity of Tmem119 immunostaining in organoid sections from 2- to 4-months. 2-month-old isogenic vs 4-month-old isogenic, p-value < 0.05. 2-month-old AD vs 4-month-old isogenic, p-value < 0.05. 2-month-old AD vs 4-month-old isogenic, p-value < 0.05. 3-month-old isogenic vs 4-month-old isogenic, p-value < 0.05. 3-month-old AD vs 4-month-old isogenic, p-value < 0.05. 3-month-old AD vs 4-month-old AD, p-value < 0.05. Data are mean +/- SD.

### Aβ localizes into large deposits in AD organoids but remains diffuse in isogenic controls

Given the amplification of glial immune responses as a result of the N1411 mutation, the dramatic down-regulation in cytokine secretion in older AD organoids rather than a persistent or worsening heightened immune response is surprising. It has been observed that under chronic Aβ exposure microglia can enter a “chronic tolerant phase” exhibiting a diminished cytokine secretion profile and impaired phagocytic ability,^43^ which can exacerbate the accumulation of proteinopathy. Correspondingly, persistent immune activation and cytokine release, which may be present at 2 months, can directly increase Aβ and tau pathology in AD. We then investigated changes in Aβ and pTau levels in the organoids that could explain the observed immune phenotype in the aging organoids (increasing Aβ overtime to dampen the immune response) and support the expected effects on pathology from the dysregulated immune signaling. Surprisingly, pTau levels of AD and isogenic control organoids did not differ from 2 to 4 months in culture (Fig 4). At 3-months, AD and control organoid immunostaining did not demonstrate any differences in Aβ or pTau signal (Fig 4A,B). There was a moderate increase in pTau levels in the 4-month organoids (Fig 4C,), but there was no difference between AD and isogenic controls.

**Figure 4:**
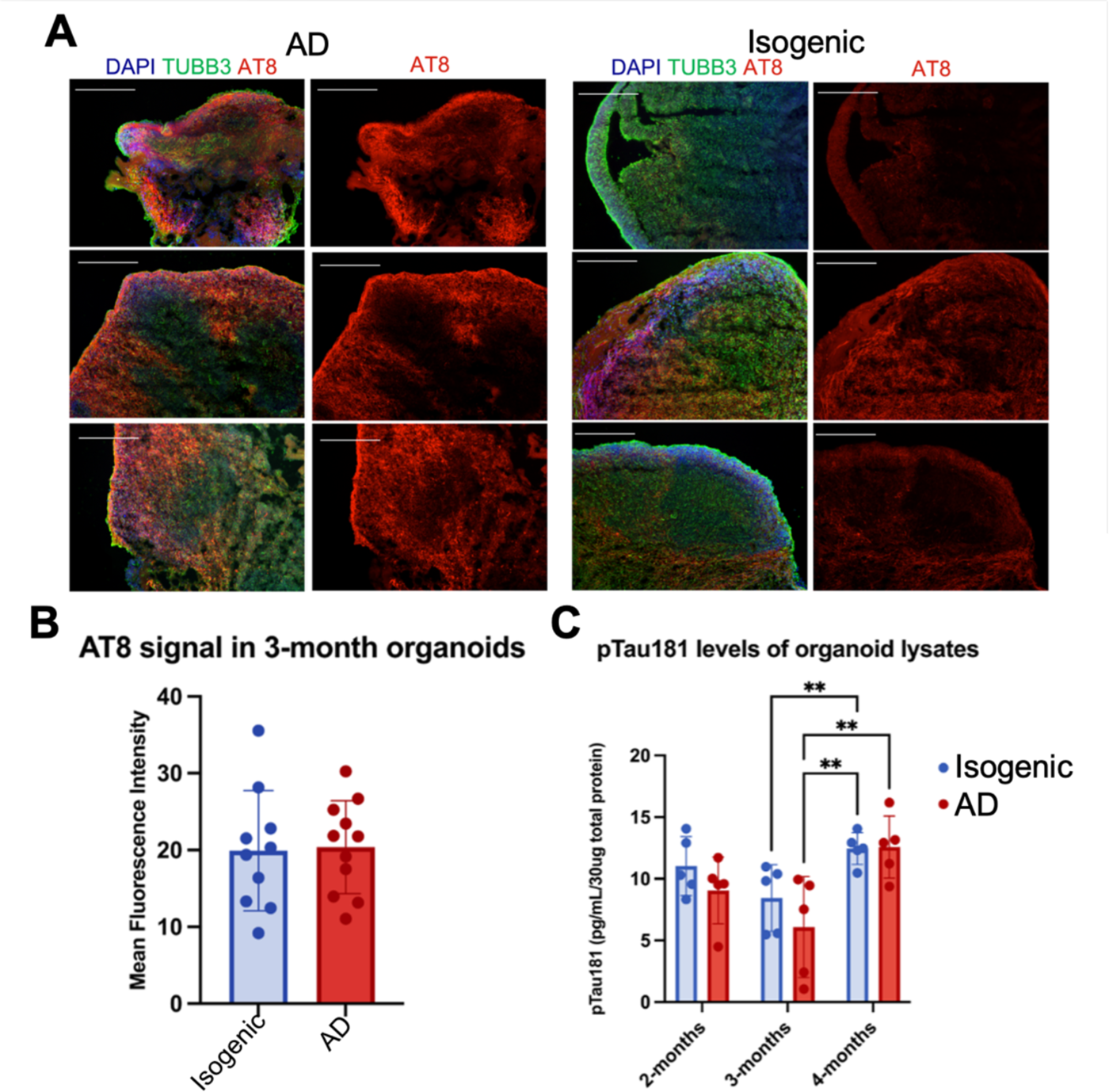
No difference in pTau levels between AD and isogenic organoids. (A) Composite (red: AT8, green: TUBB3, blue: DAPI) and red channel (AT8) images of 3-month-old AD (left) and isogenic organoids (right), scale bars: 250μm. (B) Quantification of mean fluorescence intensity of pTau AT8 immunostaining in 3-month-old organoids. Each point is a unique organoid section. Unpaired Student’s t-test. (C) pTau-181 protein levels in AD and isogenic organoid lysates from 2- to 4-months. Each sample represents the lysate of a combined 3-5 organoids. Two-way ANOVA with Tukey’s multiple comparison test. 3-month-old isogenic organoid vs. 4-month-old isogenic organoid, p-value < 0.005. 3-month-old AD organoid vs 4-month-old isogenic organoid, p-value < 0.005. 3-month-old AD organoid vs 4-month-old AD organoid, p-value < 0.005. Data are mean +/- SD.

Similarly, Aβ levels of AD and isogenic control organoids did not increase overtime in either the AD or control organoids (Fig 5). Aβ-40 and Aβ-42 levels of organoid lysates did not change over time or between AD and isogenic organoids (Fig 5A). At 3-months, AD and control organoid immunostaining similarly did not demonstrate a difference in the mean fluorescence intensity of Aβ signal (Fig 5C). However, 3-month-old AD organoids exhibited larger concentrated areas of Aβ compared to controls (Fig 5B,D). The results indicate that the AD organoids may exhibit altered deposition, rather than the accumulation, of Aβ.

**Figure 5:**
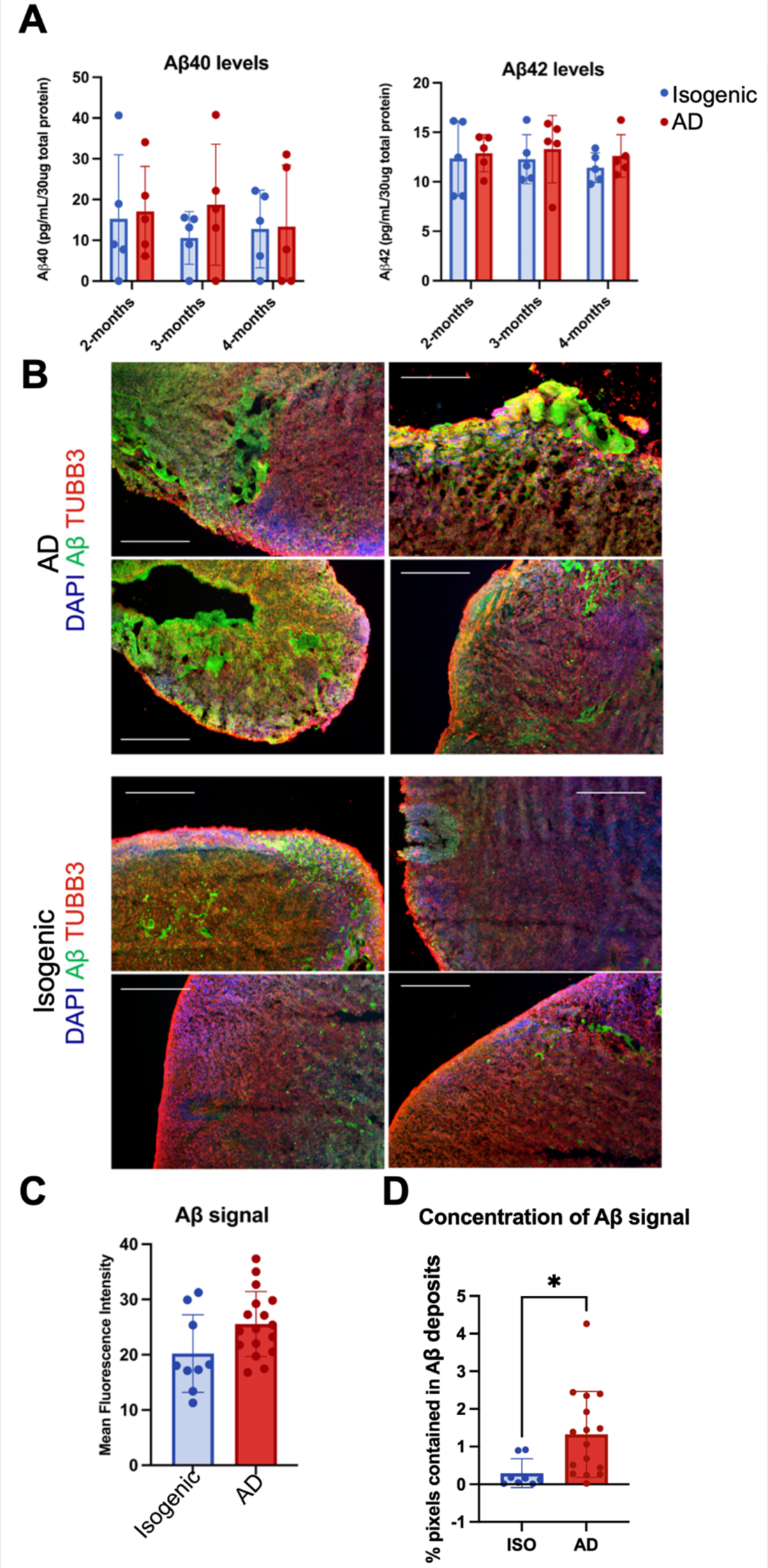
Aβ levels are constant between AD and isogenic organoids, despite more concentrated Aβ deposition in AD organoids. (A) Aβ40 (left) and Aβ42 (right) protein levels in AD organoid and isogenic organoid lysates from 2- to 4-months. Each sample represents the lysate of a combined 3-5 organoids. Two-way ANOVA with Tukey’s multiple comparison test. 3-month-old isogenic organoid vs 4-month-old isogenic organoid, p-value < 0.005. 3-month-old AD organoid vs 4-month-old isogenic organoid, p-value < 0.005. 3-month-old AD organoid vs 4-month-old AD organoid, p-value < 0.005. (B) Composite images (red: Tubb3, green: Aβ, blue: DAPI) of 3-month-old AD organoids (top) and isogenic organoids (bottom), scale bars: 250μm. (C) Quantification of mean fluorescence intensity of Aβ immunostaining in 3-month-old organoids. Each point is a unique organoid section. Unpaired Student’s t-test. (D) Quantification of Aβ deposits: percentage of pixels and neighboring pixels above threshold value (Methods). Unpaired Student’s t-test, p-value < 0.05. Data are mean +/- SD.

### Aβ levels are unchanged following microglia reduction in AD organoids, while pTau is affected from 2 to 3-months but not 2-4 or 3-4 months

To continue to tease apart a potential microglia-dependent effect on pathology in the N141I AD organoids, we investigated the depletion of microglia by colony stimulating factor 1 receptor (CSF1R) signaling inhibition with the application of the small molecule PLX5622. CSF1R signaling is required for microglia homeostasis and survival, and its inhibition results in the apoptosis of microglia.^44^ PLX5622 has been used previously to eliminate >95% of microglia *in vivo* and in *ex vivo* brain slices.^45,46^ Microglia depletion with PLX5622 has previously been studied in a variety of AD and tauopathy mouse models and often results in positive effects on tau pathology ^47–49^ but varying influence on Aβ.^45,50,51^ However, some studies demonstrate contradictory results, which may be attributed to the type of mouse model used, the presentation of pathology at the time of experimentation, and the degree of microglia depletion achieved. Microglia depletion in a human AD cerebral organoid model more reminiscent of human disease^16^ may improve our understanding of how microglia can drive AD pathology. We added 5 μM or equal amounts of vehicle control to the organoid culture medium with each medium change for the duration of the experimental timeline. PLX5622 treatment was applied to the organoids from either 2- to 3-months of age, 3- to 4-months of age, or 2- to 4-months of age to parse out the microglia-specific effects that correspond to shifting immune responses observed over time (Fig 2) and address the dynamic nature of reactive microglial states that differentially affect disease pathology in AD.^8^ Following PLX5622 treatment, we observed >50% reduction in Tmem119 signal (Fig 6A,B). The reduced Tmem119 signal intensity was not statistically different between 2- to 3-month PLX-treated AD organoids and 3- to 4-months PLX-treated AD organoids, suggesting the same level of microglial depletion was achieved from both starting points.

**Figure 6:**
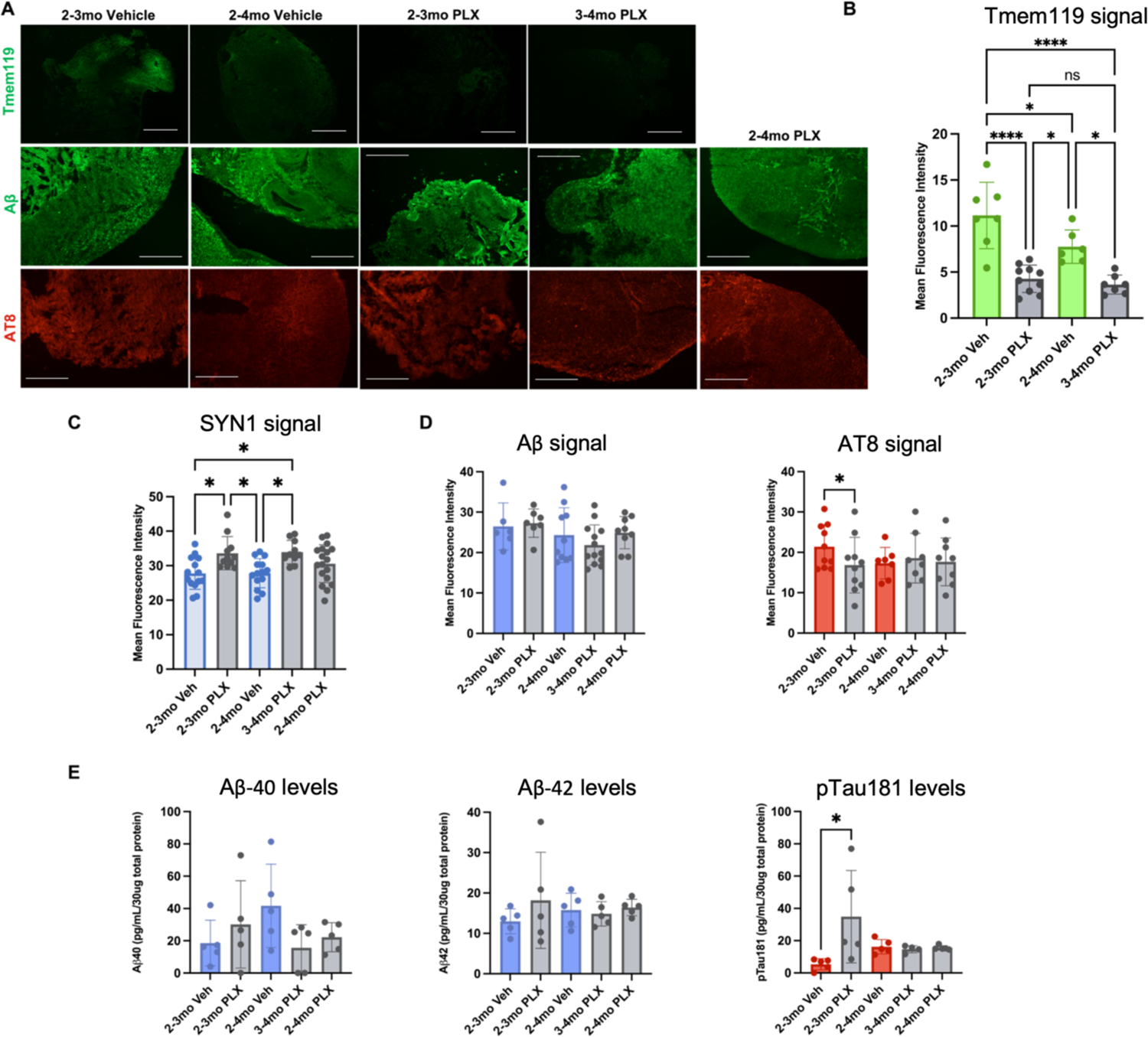
Microglia depletion prevents synapse loss, and early depletion affects tau pathology in AD organoids. (A) Representative images of microglia (Tmem119, top, scale bars: 500μm), Aβ (middle, scale bars: 250μm), and pTau (AT8, bottom, scale bars: 250μm) across PLX5622 and vehicle treatment schemes in AD organoids. (B-D) Quantification of the mean fluorescence intensity of (B) Tmem119, (C) SYN1, and (D) Aβ (left) and pTau AT8 (right) immunostaining. Each point represents the mean fluorescence intensity of one unique organoid section. Two-way ANOVA with Tukey’s multiple comparison test. Tmem119: 2- to 3-month vehicle treatment vs 2- to 4-month vehicle treatment: p-value <0.05, 2- to 3-month vehicle treatment vs 2- to 3-month PLX5622 treatment: p-value <0.0001, 2- to 3-month vehicle treatment vs 3- to 4-month PLX5622 treatment: p-value <0.0001, 2- to 4-month vehicle treatment vs 2- to 3-month PLX5622 treatment: p-value <0.05, 2- to 4-month vehicle treatment vs 3- to 4-month PLX5622 treatment: p-value <0.05. SYN1: 2- to 3-month vehicle treatment vs 2- to 3-month PLX5622 treatment: p-value <0.05. 2- to 3-month vehicle treatment vs 3- to 4-month PLX5622 treatment: p-value <0.05. 2- to 4-month vehicle treatment vs 2- to 3-month PLX5622 treatment: p-value <0.05. 2- to 4-month vehicle treatment vs 3- to 4-month PLX5622 treatment: p-value <0.05. AT8: 2- to 3-month vehicle treatment vs 2- to 3-month PLX5622 treatment: p-value <0.05. (E) Aβ40 (left), Aβ42 (middle), and pTau-181 (right) protein levels in organoid lysates. Each sample represents the lysate of a combined 3-5 organoids. Two-way ANOVA with Tukey’s multiple comparison test. 2- to 3-month vehicle treatment vs 2- to 3-month PLX5622 treatment: p-value <0.05. Data are mean +/- standard deviation.

The fluorescence intensity of Aβ and pTau staining did not change as a consequence of PLX-treatment with the exception of 2- to 3-month PLX-treatment which mildly decreased pTau AT8 levels (Fig 6A,D). Consistently, quantified levels of Aβ40 and Aβ42 in organoid lysates did not change with PLX-treatment (Fig 5E). A change in pTau-181 in AD organoid lysates was similarly only observed in the 2- to 3-month PLX-treated organoids. However, during this treatment timeline, the signal intensity of pTau-181 increased. Tau has 85 different phosphorylation sites, which variably increase the aggregation propensity of tau or decrease its association with microtubules.^52^ AT8 and pTau-181 correspond to unique phosphorylation sites on tau that are found to be increased in tauopathy. Importantly, different phosphorylation sites have been associated with specific pathology (i.e. around plaques) or location (i.e. colocalized with synapses).^53^ Similarly, numerous kinases may differentially phosphorylate tau at various sites^54^. Depending on the microglial signals that were reduced upon microglial depletion, such as a decrease in cytokine secretion that activates tau-phosphorylating kinases, the difference in affect is not unreasonable. Additionally, the semi-quantification of signal intensity does not wholly reflect protein concentration. Importantly, the sole effect on pTau occurred as a result of microglial depletion from 2- to 3-months when cytokine secretion was up-regulated (Fig 2A).

## Discussion

Neuroinflammation is not a constant, linearly increasing process in the aging diseased brain, but instead a much more convoluted series of likely co-existing cell states that are differentially responding to immune challenge. Aβ and hyperphosphorylated pathological tau both activate microglia, but there is evidence for diverse responses, such as an over-exaggerated “primed” response,^58^ a dampened, tolerant response,^43^ or even the initiation of cellular senescence.^59–63^ At the transcriptomic level, there is a great diversity of unique microglial states,^9–11^ but how these translate to their dysfunction is not completely understood. Here, we have demonstrated a dynamic immune milieu in a human cerebral organoid model as a consequence of the familial AD PSEN2 N141I mutation. The AD organoids exhibited an overall decreasing cytokine signature over months in culture compared to the isogenic controls. The AD signature uniquely reflected the underlying pathology of increased Aβ deposition and decreased synapse density. Early in AD organoid culture, increased cytokines correlated with microglia-sensitive pTau levels, as revealed by microglia depletion studies, whereas the later near-complete decrease in cytokines coincided with a decrease in synapse density that was prevented with microglial depletion. Surprisingly, levels of Aβ40 and Aβ42 did not change over time; however, Aβ immunostaining exhibited more concentrated signal as opposed to the isogenic organoids where Aβ staining was primarily diffuse.^64,65^ Microglia are thought to sequester Aβ in the form of plaques and plaque-associated microglia display an altered immune profile,^66–68^ and, as such, the increased deposition of Aβ in the AD organoids may be a result of the altered immune signaling rather than an increase in the levels of Aβ. Consistent with our findings, even in the absence of inflammatory stimuli, the microglia of N141I carrying mice demonstrate increased engulfment of Aβ.^24^ ^26^

The differences between the cytokine signatures of AD and isogenic organoids highlight key cytokines that have previously been identified in AD research. IL-4, which was up-regulated in 2-month-old AD organoids is traditionally regarded as an anti-inflammatory cytokine that can signal to microglia to adopt a phagocytic phenotype,^57^ and IL-4 treatment ameliorates AD pathology in AD mouse models.^69^ G-CSF was down-regulated in 4-month-old AD organoids. G-CSF mobilizes microglia, and, consistently, microglia in AD exhibit impaired motility.^70,71^ G-CSF treatment has been shown to rescue cognitive impairment in AD mouse models.^72,73^ Fractalkine, or CX3CL1, is a transmembrane protein that is constitutively expressed by neurons.^74^ Temporal recruitment of microglia by fractalkine signaling directs homeostatic synapse pruning and network maturation,^75^ and injured neurons also cleave fractalkine from their cell membrane to recruit microglia, resulting in their death.^76^ Fractalkine was decreased in 4-month-old AD organoids but increased at earlier time points. GRO (CXCL1), up-regulated in early AD organoids, has been linked to the phosphorylation of tau and the production of reactive oxygen species.^77,78^ 2- and 3-month AD organoids also exhibited increased FGF-2, which was subsequently down-regulated at 4-months. FGF-2 is secreted by injured neurons to initiate protective microglial phenotypes.^79^ Most notably, the PLS-DA models showed key contribution of the up-regulation of VEGF in the AD organoids at 3-months and the sole up-regulation of VEGF in 4-month-old AD organoids. VEGF has previously been identified through PLS modeling in Alzheimer’s disease patient data by our group and was demonstrated to decrease neuron viability in the presence of Aβ.^80^ The makeup of the cytokine signatures convey a complicated depiction of the overall immune state with simultaneous protective and detrimental activities; however, it is correlated with a larger picture of decreasing synapse density and accumulation of concentrated Aβ.

Synapse loss is one of the earliest alterations in AD and is closely associated with microglia activation in human imaging studies.^81–83^ AD mouse models have demonstrated microglia’s involvement thorough their direct engulfment of synapses,^84,85^ and synapse loss is observed before plaque deposition.^86,87^ The prevention of synapse loss in AD organoids with the depletion of microglia, support their direct role in aberrant synapse engulfment, and, importantly, this occurred prior to the chronic immune insult of substantial pathology accumulation. Although Tmem119 signal intensity increased in both the AD and isogenic organoids, suggesting an increasing microglial population, only increasing Tmem119 signal in the AD organoids coincided with decreasing synaptic density, implicating the dysregulated immune response of AD organoids. The very mild, if any, change in the Aβ and pTau levels over time in the organoids is quite remarkable, given the known numerous microglial drivers of Aβ and pTau pathology.^3^ This could suggest that the microglia are driving the observed changes, i.e Aβ deposition and synapse loss, independently of pathology. Others have also suggested a larger role of microglia in neurodegeneration, including one study that determined microglia-mediated neuronal damage, rather than tau-dependent mechanisms, was the primary contributor of neurodegeneration in a tauopathy mouse model.^47^ It has also been shown that activated microglia alone can initiate neurodegeneration.^6^

It has been demonstrated that the N141I mutation results in the exaggeration of the glial immune response, which resembles a “primed” phenotype commonly observed in age and with chronic Aβ insult.^27^ This heightened immune response is largely observable after immune challenge, where Aβ-stimulation or LPS treatment causes an increase in the cytokine secretion of N141I-expressing glial cells compared to controls.^24,25^ We did not observe an increase of Aβ over the experimental timeline but observed an amplified immune milieu in 2-month-old AD organoids. With the normal accumulation of Aβ in age and insult,^88,89^ the early observed phenotypes from the N141I mutation may evolve over time. The events observed over the experimental timeline may be more reminiscent of early events in disease and could inform how glial dysfunction would contribute to the eventual development of AD.

Following treatment with 5μM PLX5622, we observed more than 50% reduction in Tmem119 signal. It has been previously demonstrated that 1uM PLX5622 can achieve nearly 99% reduction of microglia in organotypic slice culture.^90^ This discrepancy may be due to the size of the organoids if the compound cannot penetrate far enough into the organoid to deplete persistent microglia populations. Also, because of the immaturity of the cells and heterogeneous expression of microglia specific genes including CSF1R,^22^ the cells may not be as sensitive to PLX5622 treatment. However, in our organoids, increasing the concentration to 10μM and 20μM PLX5622 resulted in the organoids falling apart, although concentrations less than 20μM did not previously demonstrate negative effects on viability.^90^ Other strategies may have to be employed to achieve full microglia knockout in the organoids, such as a drug antibody conjugates. It is worth noting that we used the expression of Tmem119 to quantify the reduction in the microglia population. Tmem119, thought to be involved in the regulation of the microglial immune response, is differentially expressed in homeostatic and reactive states.^37^ Thus, a reduction in the associated fluorescence intensity does not directly correspond to an equal reduction in the microglial population, though this would also be the case of IBA1 as it is regarded as a marker of microglial activation.

In our study, the AD organoids (and isogenic controls) were derived from a single cell line from a male individual. In contrast to our AD organoids’ longitudinal reduction of cytokine secretion, previous studies have shown iPSC-derived microglia carrying the N141I mutation exhibit increased secretion of pro-inflammatory cytokines; however, the N141I-carrying iPSCs were derived from females.^26^ We differentiated organoids from a single cell line from a female AD patient to investigation the potential for a more “canonical” neuroinflammatory profile. The AD organoids derived from the female cell line secreted a larger diversity of cytokines and PLS regression demonstrated persistent increase of select cytokines overtime. The organoids did not exhibit decreased synapse density between 2- and 4-months in culture (Supplementary Figure 4).. While it is appreciated that there is a greater degree of neuroinflammation and neuropathology in females,^64,65^ it is unclear whether the difference between the AD organoids can be attributed to biological sex or inter-individual variation.

Further investigation with organoids derived from more iPSC lines from unique male and female individuals is needed. ^27^With the use of genetically modifiable human cells, the organoid model will be particularly critical in the discovery of the unique roles of AD-specific mutations and risk factors in early disease events through the activities of specific cell-types and their cell-to-cell communication.

We have previously identified an Aβ-specific cytokine signature in 5xFAD mice that reduces neuronal mitochondrial metabolism, capable of predisposing neurons to injury in the absence of Aβ and pTau pathology.^32^ In this study, the dynamic cytokine signature of human organoids corresponds to reduced synaptic density and the changes to the deposition of Aβ. Our studies and those of others are highlighting the importance of early glial involvement that may drive neuronal injury to a greater extent than the accumulation of proteinopathy.^6,91^ However, preventing dysfunctional immune responses is made more complicated by their diversity and dynamic nature. In this study, over a relatively short culture period, there were numerous dynamic changes in AD organoids, including variable cytokine regulation, the initial increase and subsequent decrease of synaptic density, and microglia-sensitive changes in pathology. Over the course of human disease spanning decades, the magnitude and diversity of changes is imaginably quite impressive, and further investigation will be needed to determine what initiates these changes and when and how is best to modulate them to ameliorate disease.

## Conclusion

In this study of human iPSC-derived Alzheimer’s disease cerebral organoids carrying the PSEN2 mutation N141I, we demonstrated dynamic immune signatures that corresponded to underlying changes in pathology. The AD organoids demonstrated a dampened cytokine secretion profile over time that uniquely coincided with synapse loss. The decreased synapse density in the AD organoids was prevented with microglial depletion. While absolute levels of Aβ and pTau did not increase, Aβ deposition was highly localized in the AD organoids as compared to isogenic control organoids, which demonstrated diffuse Aβ signal. Through the use of human patient-derived models, gene-specific changes capable of driving disease will identify early disease-causing mechanisms, likely unobservable in mouse model.

## Acknowledgements

This work was supported by start-up funds from the Penn State College of Medicine Departments of Neurosurgery and Pharmacology (EAP) and a grant from the H.G. Barsumian MD Memorial Fund. MKK was supported by training fellowship T32NS115667 from the National Institute of Neurological Disorders and Stroke. The authors thank Dr. Amanda Snyder for her help troubleshooting immunohistochemistry of organoid tissue, Dr. Rebecca Fleeman for help in organoid culture maintenance, Stem Cell Tech for their guidance and troubleshooting with stem cell culture, and Dr. Madeline Lancaster for her guidance in optimizing the protocol for generating cerebral organoids in our laboratory.

## Notes

### Competing Interest Statement

The authors have declared no competing interest.

### Summary of Updates

Addition of single-cell RNA sequencing and three co-authors contributing to performing this experiment and analysis.

