## Supplementary Materials for "Dynamic neuroinflammatory profiles predict Alzheimer’s disease pathology in microglia-containing cerebral organoids"

### Supplementary Methods

**Differentiation and analysis of cerebral organoids derived from female iPSC line:** The iPSC line carrying the N141I PSEN2 mutation (Coriell Institute AG25370, white female, 81 years-old at sampling) derived from a female familial AD patient was used to generate cerebral organoids and were collected as describes in "Methods." Cytokine concentrations were quantified on the Luminex FLEXMAP3D platform using the Milliplex human cytokine/ chemokine/ growth factor panel A magnetic bead panel kit (HCYTA-60K, Millipore). For PLS modeling the proteins IL-17E/IL-25, IL-17F, IL-18, IL-22, IL-27, M-CSF, MIG/CXCL9, PDGF-AA, PDGF-AB/BB were removed from the dataset, so the male and female organoid cytokine datasets consisted of the same cytokines for multivariate analysis. PLS regression (PLSR) models were generated for the prediction of continuous numerical outcomes. Repeated random sub-sampling cross-validation (repeated 100 times with 1/5$^{th}$ of the data left out for each test set (n=15)) was used to estimate the average root mean squared error of cross validation (RMSECV) to determine the optimal number of LVs for each model. Models containing 1 LV are visualized using 2 LVs, but only 1 LV was used in the calculation of the model's RMSECV and significance.

**Supplementary Figure 1**

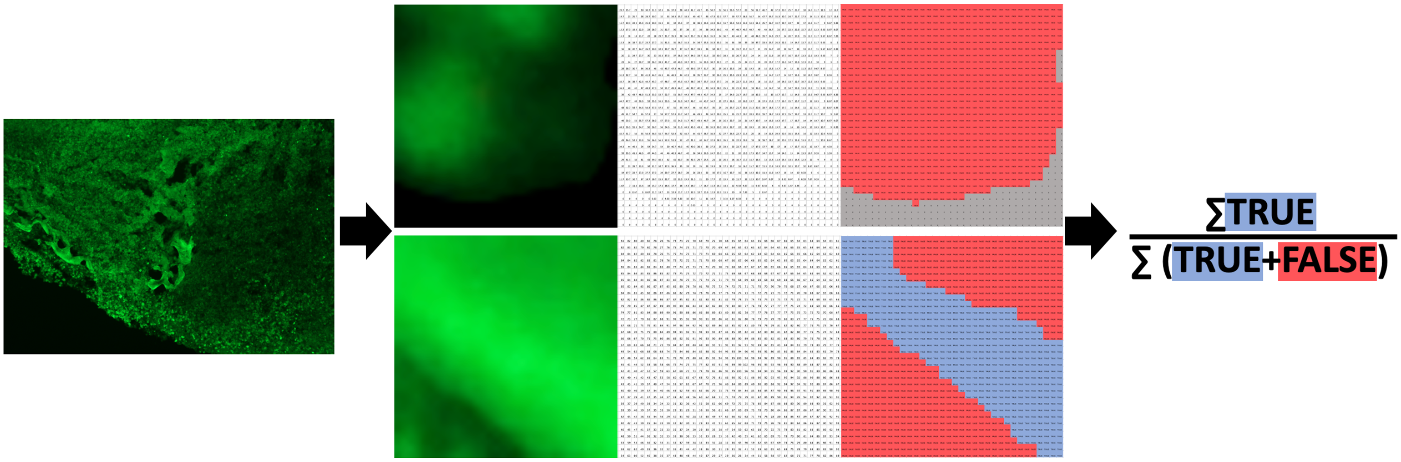


**Schematic demonstrating Aβ signal deposition quantification workflow.** For the quantification of the concentration of Aβ immunostaining, the background was deleted from each image and the image was converted into a text file indicating the fluorescence intensity of each pixel. Each pixel above the chosen threshold value was denoted as “TRUE” if all of its neighboring pixels were also above the threshold value. Otherwise the pixel would be denoted as “FALSE”. The percentage of TRUE pixels over the total number of pixels was calculated for each image (background pixel were excluded) to represent the portion of the organoid section with a strong concentrated Aβ signal. The chosen threshold value (Methods) is stringent and does not include all small regions of concentrated Aβ signal.

### Supplementary Figure 2

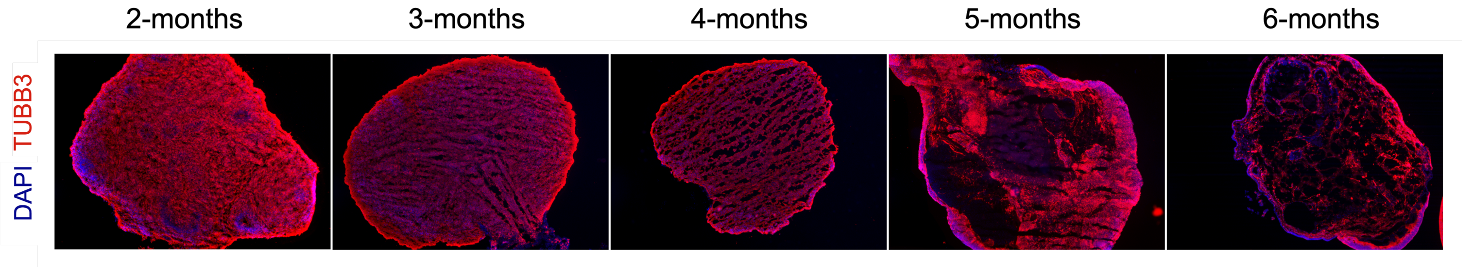


**Female PSEN2 N141I organoids exhibit robust tissue atrophy after 5-months in culture.** TUBB3 (red) and DAPI (blue) immunostaining of 2-, 3-, 4-, 5-, and 6-month-old Female PSEN2 N141I organoids (left to right).

**Supplementary Figure 3**

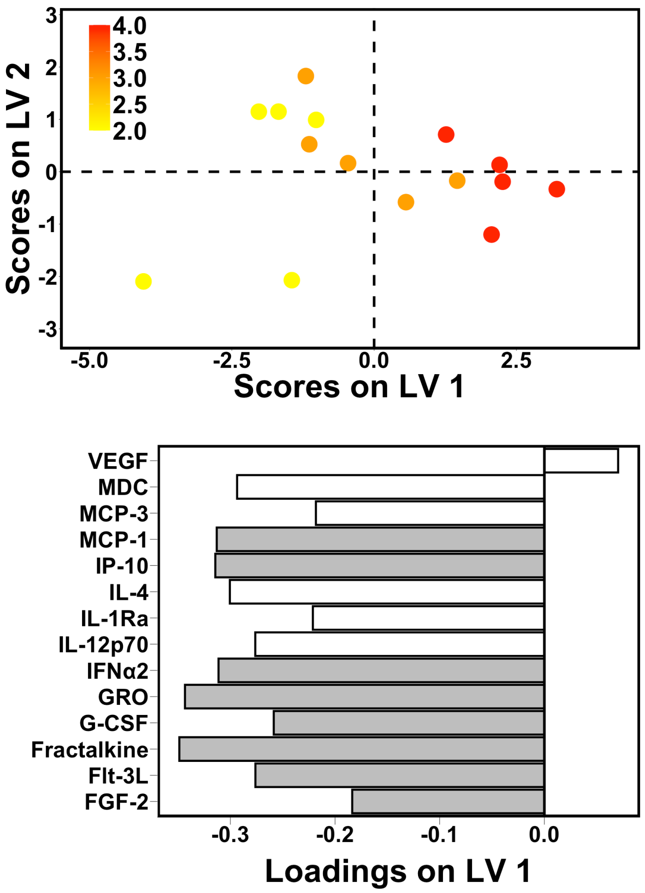


**Cytokine secretion decreases over time in AD organoids derived from the male N141I iPSC line.** PLSR scores plot (top) and LV1 loadings plot (bottom) of AD organoids (male N141I iPSCs) from 2- to 4-months in culture (1 LV, RMSECV: 0.745, p-value: 0.01) Shaded loadings indicate a VIP score > 1. Each point represents a single sample comprising the lysates of 3-5 organoids. Samples with positive scores (aged AD organoids) correlate with increased levels of cytokines with positive loadings and the down-regulation of cytokines with negative loadings.

### Supplementary Figure 4

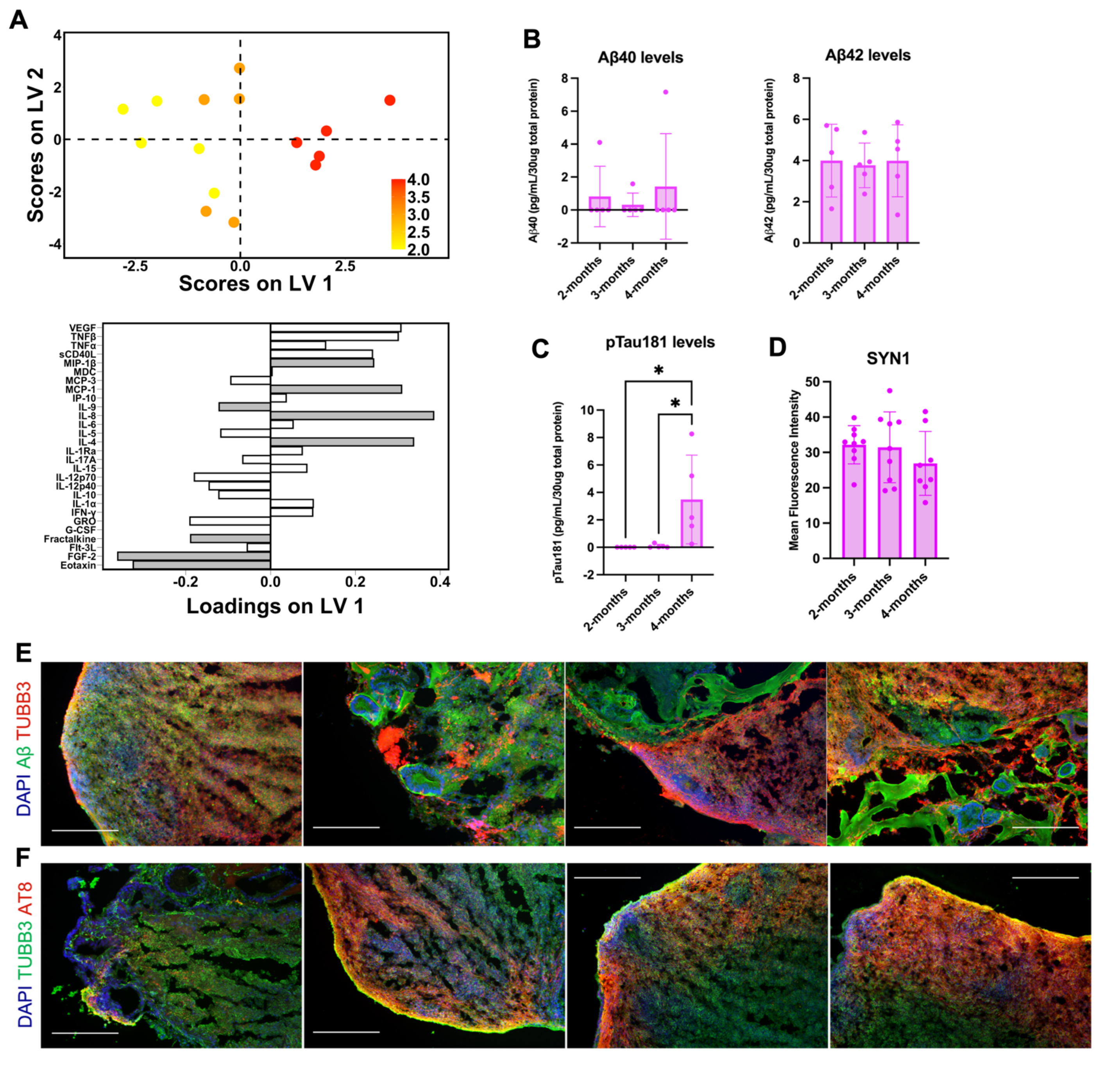


**Longitudinal cytokine secretion and pathology of AD cerebral organoids derived from the female N141I iPSC line.** (A) PLSR scores plot (top) and LV1 loadings plot (bottom) of female AD organoids from 2- to 4-months in culture (1 LV, RMSECV: 0.766, p-value: 0.01). Shaded loadings indicate a VIP score > 1. Each point represents a single sample comprising the lysates of 3-5 organoids. Samples with positive scores (aged AD organoids) correlate with increased levels of cytokines with positive loadings and the down-regulation of cytokines with negative loadings. (B) Aβ40 (left) and Aβ42 (right) protein levels in female organoid lysates from 2- to 4-months. Each point represents a lysate sample of a combined 3-5 organoids. One-way ANOVA with Tukey's multiple comparison test. (C) pTau181 protein levels in female organoid lysates from 2- to 4-months. Each point represents a lysate sample of a combined 3-5 organoids. One-way ANOVA with Tukey's multiple comparison test. 2-month-old vs 4-month-old organoids, p-value < 0.05. 3-month-old vs 4-month-old organoids, p-value < 0.05. (D) Quantified SYN1 fluorescence intensity. One-way ANOVA with Tukey's multiple comparison test. (B-D) Data are mean +/- SD. (E) Composite images (red: TUBB3, green: Aβ, blue: DAPI) of 3-month-old organoids, scale bars: 250μm. (F) Composite images (red: AT8, green: TUBB3, blue: DAPI) of 3-month-old organoids, scale bars: 250μm.
